# Integration of ‘what’ and ‘when’ predictions during action-effect processing

**DOI:** 10.64898/2026.07.27.740928

**Authors:** Wai Ying Chung, Álvaro Darriba, Florian Waszak

## Abstract

Voluntary actions generate predictions about their sensory consequences, but it remains unknown whether predictions arising from different dimensions of intentional action are processed independently or integrated into a unified expectation. We addressed this question by combining action choice (“what”) and action timing (“when”) within a single action-effect learning paradigm. Participants learned independent associations between each action dimension and a distinct auditory feature and performed self-paced actions while electroencephalography was recorded. This design allowed action-choice and action-timing predictions to be fulfilled or violated independently or simultaneously. Event-related potentials were analysed using Bayesian linear mixed-effects models. The P2 component showed a significant interaction between action-choice and action-timing prediction violations: neither violation alone reliably modulated P2 amplitude, whereas simultaneous violations produced a marked reduction, consistent with a conjunctive prediction-error response. The later P3 component exhibited a graded pattern, with action-timing violations increasing amplitude and action-choice violations contributing an additional increment when both predictions were violated. These findings suggest that predictions derived from different dimensions of voluntary action are integrated during early sensory processing, whereas later stages evaluate prediction violations more cumulatively. The present results extend previous work on action-effect prediction by showing that predictions associated with action choice and action timing interact during the processing of their sensory consequences.

## Introduction

To interact adaptively with our environment, human behaviour must be fundamentally goal-directed. Goal-directed behaviour requires a deep, learned knowledge of the relationship between actions and their consequences. Accordingly, the classic ideomotor principle assumes that actions are selected, planned, and executed based on the active anticipation of their sensory consequences, such that the expectation of a desired sensory outcome is an integral part of action planning rather than a consequence of movement alone (Greenwald, 1970; Prinz, 1997). Extending this idea, the common coding theory proposes that actions and their perceptual effects share common neural representations, allowing anticipated sensory consequences to directly drive action selection, planning, and execution (Hommel, 2019; Prinz, 1990). Through experience, these action-effect associations enable the brain to predict the sensory consequences of self-generated actions, supporting efficient interaction with the environment, adaptive control, and the sense of agency (Blakemore et al., 1998; Wolpert & Flanagan, 2001).

According to predictive coding frameworks, the brain implements these action-effect associations by continuously comparing top-down sensory predictions with bottom-up sensory inputs. Any mismatch between the expected and actual input is computed as a prediction error (PE), which serves to update the internal predictive model and minimise future errors through iterative refinement (Friston, 2005). In electrophysiological experiments on action-effect prediction, this PE signal is commonly operationalised using event-related potentials (ERPs). A robust finding is that self-generated actions, whose sensory consequences are highly predictable, elicit attenuated amplitudes of the auditory N1 and, in some cases, the P2 components compared to externally generated or involuntary actions (Bendixen et al., 2012; Horváth, 2015; Timm et al., 2014). This attenuation is interpreted as a reduction in PE, reflecting the successful anticipation of the action’s sensory outcome. Other studies have employed active oddball paradigms in which participants perform intentional actions that sometimes produce unexpected sensory effects (Hughes et al., 2013; Korka et al., 2019). In this type of design, predicted action-effects typically evoke smaller N1 amplitudes (Hughes et al., 2013; Hughes & Waszak, 2011), while mispredicted effects elicit larger P2 amplitudes (Korka et al., 2019). This P2 modulation has been linked to the mismatch negativity (MMN), a well-established ERP index of auditory PE and predictive model updating (Garrido et al., 2009; Wong et al., 2024). Prediction violations have also been associated with modulations of the later P3 component, which is generally thought to reflect the evaluation of unexpected events and the updating of internal models in response to behaviourally relevant PE (Mars et al., 2008; Polich, 2007). Together, the P2 and P3 provide complementary indices of the neural processes underlying the detection and evaluation of action-effect PE.

Following the ideomotor account, action-effect prediction has traditionally been studied as a unitary process. However, voluntary actions themselves are not unitary. Brass and Haggard (Brass & Haggard, 2008) proposed that intentional action comprises three partially dissociable decision components: deciding what action to perform, when to perform it, and whether to perform it at all. Neuroimaging studies have provided evidence that these components recruit partly distinct neural networks, suggesting that different aspects of voluntary decision-making may contribute independently to action planning and control (Krieghoff et al., 2009; Mueller et al., 2007; Zapparoli et al., 2017).

Building on this framework, Chung et al. (Chung et al., 2022) investigated whether predictions about action consequences are generated from each of these intentional components. Using an action-effect learning paradigm, they showed that violations of expected auditory consequences elicited comparable prediction-related ERP responses regardless of whether predictions were based on action choice (“what”), action timing (“when”), or action execution (“whether”). Their findings demonstrated that action-effect prediction is a general property of intentional action rather than being specific to a particular decision dimension.

Although this work established that different dimensions of voluntary action can independently generate sensory predictions, the study considered each dimension in isolation. In everyday behaviour, however, actions rarely involve independent decisions about what or when to act. Instead, a single voluntary action simultaneously specifies multiple characteristics of its expected sensory consequences. For example, striking a piano key determines both which sound will be produced and when it will occur. Consequently, the brain must combine predictions arising from multiple dimensions of action into a coherent expectation of the forthcoming sensory event.

Recent evidence from perceptual predictive processing suggests that such integration may indeed occur. Using auditory sequences in which stimulus identity (“what”) and temporal regularity (“when”) were manipulated independently, a recent study (Cappotto et al., 2023) showed that PEs for sound identity depended on temporal expectations, indicating that predictions about what will occur and when it will occur are integrated rather than processed independently. Importantly, these findings were obtained for externally generated auditory events, where predictions arise from learned sensory regularities rather than from voluntary action.

Whether the same principle applies to action-effect prediction remains unknown. Predictions derived from action choice and action timing originate from internally generated motor decisions rather than external sensory statistics and may therefore be combined differently from sensory-based predictions. The present study addressed this question by combining action choice (“what”) and action timing (“when”) within a single action-effect learning paradigm. Participants learned independent associations between each action dimension and a distinct auditory feature, allowing predictions based on action choice and action timing to be fulfilled or violated either independently or simultaneously. This factorial design enabled us to determine whether neural responses to unexpected action outcomes reflect separate PEs for each action dimension or a unified representation of the expected sensory consequence.

Two competing accounts can be considered. If action-choice and action-timing predictions are represented independently, violations of each prediction should produce separable effects on neural responses, with simultaneous violations resulting in approximately additive PE signals. Alternatively, if the brain integrates multiple action-derived predictions into a single representation of the expected sensory event, simultaneous violations should produce a non-additive PE response because the predicted sensory event as a whole has been violated.

To test these alternatives, we recorded EEG while participants performed the task and analysed the auditory N1, P2, and P3 ERP components, which have previously been implicated in the detection and evaluation of action-effect PE. Rather than predicting a specific pattern of ERP modulation, our primary aim was to determine whether neural responses to unexpected action outcomes reflect independent contributions of action-choice and action-timing predictions or evidence that these predictions are integrated into a unified representation of the expected sensory consequence. Because previous studies have established that each action dimension can independently generate sensory predictions (Chung et al., 2022), but have not examined how multiple action-derived predictions interact, both outcomes were theoretically plausible. Demonstrating independent effects would suggest that PEs are computed separately for different dimensions of voluntary action, whereas evidence for integration would indicate that the motor system combines multiple action-derived predictions into a coherent sensory expectation. More broadly, establishing such integration would extend previous work on action-effect prediction (Chung et al., 2022) and provide evidence that the combination of *what* and *when* information represents a general principle of predictive processing, operating not only during perception of externally generated events (Cappotto et al., 2023) but also in predicting the sensory consequences of one’s own actions.

## Methods

### Participants

Participants gave written informed consent, and experimental procedures were undertaken in accordance with the Declaration of Helsinki and with the approval by the Comité de Protection des Personnes Ile de France II. Participants received monetary compensation for their participation. All participants reported normal hearing and normal or corrected-to-normal vision, none reported any history of neurological conditions. Data from one participant were excluded due to incomplete data, and data from three participants were excluded due to an excessive number of noisy trials (> 15% of trials rejected). The final sample consisted of 25 participants (14 females, 11 males; 3 left-handed; mean age = 27.9 years, age range = 18–39 years). Previous EEG studies on auditory processing and PE signals have found significant effects on event-related potentials, namely N1, P2, and P3, with similar or smaller sample sizes (Hsu, Bars, et al., 2015; Korka et al., 2019; Timm et al., 2014).

### Stimuli and procedures

Auditory stimuli consisted of four tones differing in pitch and waveform. Two tones were generated using a triangle waveform at the frequency of D4 (293.66 Hz) and A4 (440.00 Hz), and two tones were generated using a sawtooth waveform at the same frequency (D4 and A4). All tones had a duration of 150 ms (including 6 ms rise/fall times) and were presented binaurally at a fixed volume through a headphone. Tones were generated digitally at a sampling rate of 44100 Hz.

All stimuli were presented at the centre of the screen against a grey background on a 27-in., 60 Hz LCD display, 100 cm from the participants. The task was delivered with Psychotoolbox-3 (Kleiner et al., 2007) running on MATLAB. Prior to the experiment, participants received both written and verbal instructions that explained the procedure of the experiment and performed 240 practice trials. Participants were instructed to place their index fingers on the “S” and “L” keys on the keyboard.

The procedure of a single trial is illustrated in Fig. 1. Each trial began with a filled black square (200 × 200 pixels) centrally positioned on a grey screen. The square then enlarged from the centre outward, reaching its maximum size (400 × 400 pixels) over approximately 500 ms, before shrinking back to its original size (200 × 200 pixels) over the same duration. Participants were instructed to press either a left key (S) or a right key (L) at a time of their own choosing. A response made during the enlarging phase (100–500 ms from stimulus onset) was classified as an early response, and a response made during the shrinking phase (600–1000 ms from stimulus onset) was classified as a late response. Responses made within 100 ms of stimulus onset were flagged as too quick and a warning message was displayed. Trials on which no response was recorded were not counted and the trial was repeated.

**Figure 1.**
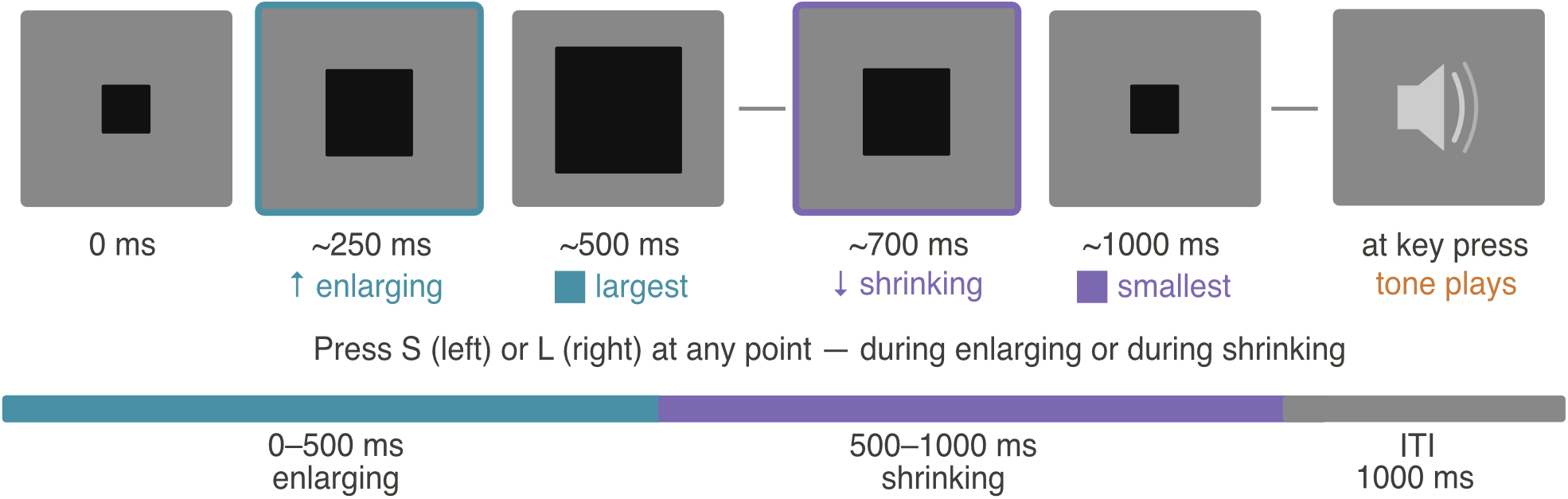
Schematic representation of a single trial. On each trial, participants freely chose which key to press (left or right) and when to press it (during the enlarging or shrinking phase of the visual stimulus). The keypress triggered a tone whose pitch and waveform were predicted by the choice of key and the timing of the response, respectively.

The tone presented following each key press was determined by the combination of action choice (left or right key) and action timing (early or late response). Tone frequency (293.66 Hz or 440.00 Hz) and waveform (triangle or sawtooth) were each mapped onto one of the two action dimensions, such that one dimension predicted tone frequency and the other predicted tone waveform. For each mapping, the predicted tone occurred with 80% probability and the alternative tone occurred with 20% probability. For example, if action choice was mapped onto frequency and the right key was associated with the high-frequency tone (440.00 Hz), pressing the right key would trigger a high-frequency tone on 80% of trials and a low-frequency tone on the remaining 20%. Whether action choice predicted frequency or waveform, and correspondingly whether action timing predicted waveform or frequency, was counterbalanced across participants. Additionally, the mapping of specific keys and timing phases onto the two levels of their respective predicted tone properties (high /low frequency, or triangle/sawtooth waveform) was also counterbalanced across participants.

The experiment consisted of 2400 trials in total, with self-paced breaks every 100 trials. Participants were required to make at least 600 responses for each combination of action choice and action timing and were shown the remaining count for each combination at every break to encourage balanced sampling across the four action combinations.

### EEG data acquisition and preprocessing

We recorded the EEG using the PyCorder system and actiCHamp amplifiers (BrainProducts GmbH, Gilching, Germany) in DC recording mode with a sampling rate of 2000 Hz. Continuous EEG data were collected from 60 actiCAP EEG electrodes (BrainProducts GmbH) mounted on an elastic cap and referenced to the right mastoid. EEG electrodes were arranged following the extended 10–10 position system (Acharya et al., 2016). Additional electrodes were placed on the right/ left mastoid and on the outer canthi of both eyes. Custom-built MATLAB scripts with EEGLAB (Delorme & Makeig, 2004) functions were used for the pre-processing of the EEG data. EEG data was filtered offline (high pass: 0.1 Hz, low pass: 40 Hz) and re-referenced to linked mastoids. Bad channels were identified by visual inspection of the EEG raw data and the channels ’power spectra. Independent component analysis (ICA) was performed to identify components that were associated with blinks and eye movements. Components containing those artifacts were rejected by visual inspection and measures computed with the EEGLAB plug-in functions IClabel (Pion-Tonachini et al., 2019). Subsequently, we removed all trials in which activity exceeded ±100 μV to account for noise and large muscle artefacts, resulting in the average exclusion rate of 2.6% (SD = 5.12). Bad channels were reintroduced by interpolating data between neighbouring electrodes using spherical spline interpolation (Perrin et al., 1987). Epochs were extracted from -200 ms to 1000 ms relative to stimuli onset and baselines were corrected to the 200 ms pre-stimulus interval. To minimise the influence of individual differences in scalp topographies and to reduce the impact of multiple statistical comparisons, ERP components were analysed using regions of interest (ROIs) defined by relevant electrode sites. These ROIs were selected based on both the grand-average visual identification of peak electrodes and the topographical distribution of scalp activity (see Fig. 2). Following this procedure, the N1 component was measured in the latency range of 70–100 ms at FCz, while P2 and P3 were measured in the latency ranges of 150–180 ms and 300–350 ms, respectively, at FCz and Cz. The chosen time windows and electrode sites for the N1, P2, and P3 components are consistent with previous literature (Bäß et al., 2008; Darriba & Waszak, 2018; Hsu, Bars, et al., 2015; Hughes et al., 2013).

**Figure 2.**
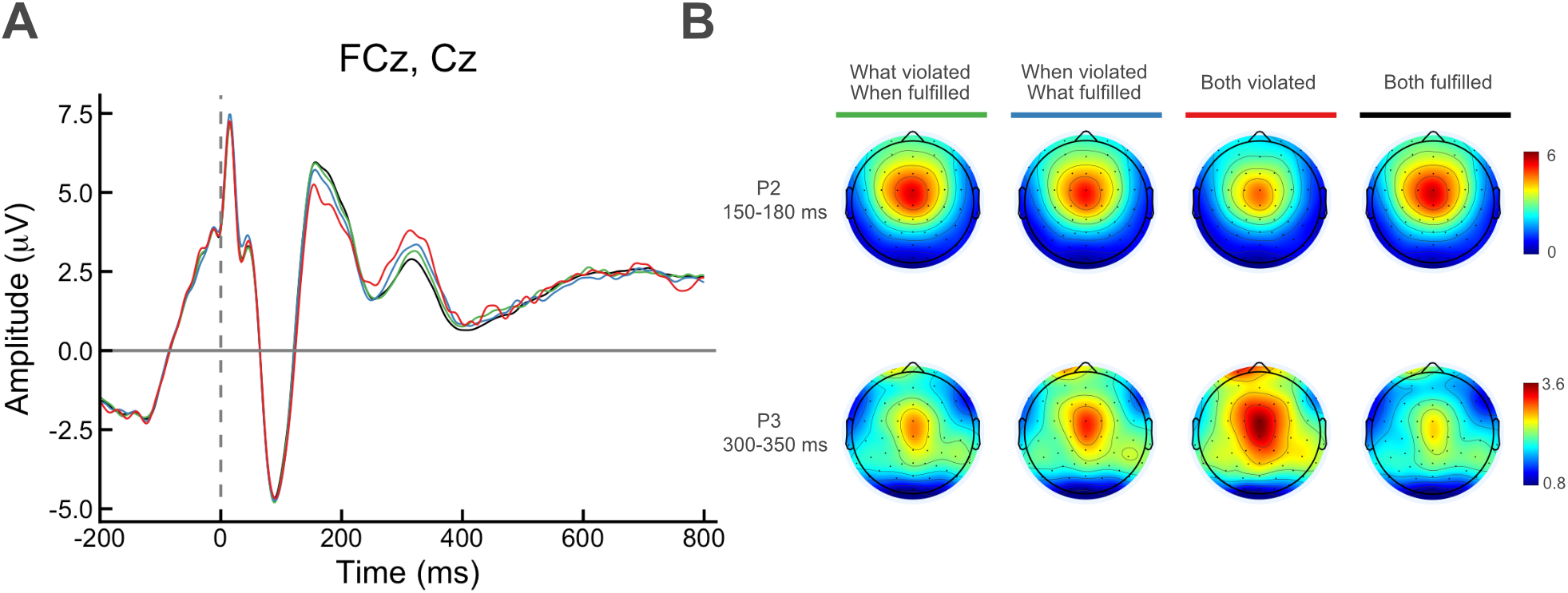
(A) ERP waveforms corresponding to the average of FCz and Cz electrodes, showing the N1 (70–100 ms), P2 (150–180 ms) and P3 (300–350 ms) components. (B) Topographic maps of average amplitude in P2 and P3 time window for every condition.

### Data analysis

We investigated ERP correlates of auditory prediction by comparing responses across four conditions defined by whether the prediction associated with action-choice (what-prediction) and action timing (when-prediction) were fulfilled or violated, either individually or in combination. Single trial EEG data were analysed with a Bayesian linear mixed-model (LMM) analysis using the package brms (Bürkner, 2017), a high-level interface on Stan (Carpenter et al., 2017) in R. Plots were made using brms and ggplot2 (Wickham, 2016). An advantage of LMMs over traditional approaches such as repeated measures ANOVA and paired sample t-tests is that a single model can take all sources of variance into account simultaneously. Furthermore, comparisons between conditions can easily be implemented in a single model. LMMs (of which t-tests and ANOVA are specific examples) allow for modelling complex data structures and taking correlations in data structures into account. Bayesian LMMs do so in a more powerful way than maximum likelihood models, even with small sample sizes. With weakly informative priors, Bayesian analysis gives insight into the range of possible effect sizes, reduces possible overinterpretation of sampling error, and allows for direct comparison of effect sizes. It is theoretically distinct from frequentist statistics in its inferences. The coefficient estimates are expressed in credible intervals. Credible intervals reflect the intuitive notion of the value of a parameter falling within that interval with a given probability, 95% in this case.

We used a predefined model reflecting our experimental design (Barr et al., 2013), and we kept this model structure the same across ERP components. Participant amplitudes were normally distributed and did not need transformation to their logarithmic function (Baayen, 2008). In the model, observations were predicted by Action choice/What-prediction (violated vs. fulfilled) and Action-timing/When-prediction (violated vs. fulfilled) in a full interaction. The model additionally included individual participant intercept as random effect and predictors as random slopes to account for individual variation regarding the experimental effects. Factorial predictors were treatment-coded with the fulfilled level as the reference category (coded 0) and the violated level as the comparison (coded 1). Under this coding, the model intercept represents the estimated amplitude in the Both fulfilled condition, the coefficients for what-violation and when-violation represent the difference from Both fulfilled when each violation occurs in isolation, and the interaction coefficient captures any additional amplitude difference when both violations co-occur beyond the sum of their individual effects. Planned pairwise comparisons were conducted via Bayesian hypothesis testing using the function Hypothesis in brms with Bonferroni correction. We used a generic weakly informative prior with mean 0 and 1 SD over the fixed effects and kept all other priors at default. We used 4 chains of 3000 iterations each per model, of which 1000 per chain were used for warm-up only, a maximum tree depth of 15 and a target acceptance rate (adapt delta) of 0.95. Convergence was verified through visual inspection of trace plots, and the Rhat of 1.00 for each parameter.

The model was specified as follows,

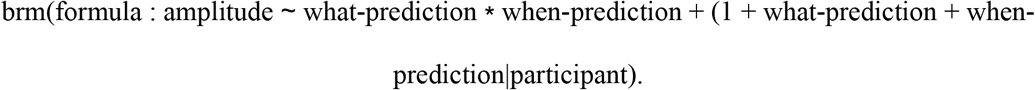

## Results

### Behavioural performance

Participants generally performed well and followed task instructions. Invalid responses caused by failing to press a key within the trial response window or before the presentation of the visual stimulus were rare, averaging 3.34% (SD = 3.39%) of trials. The mean of the maximum number of consecutively repeated responses was 11.6 (SD = 8.82).

### ERP results

The results obtained in the statistical analyses are graphically and numerically illustrated in Figs.3. Since no significant effect was observed for the N1 component, the reported results are limited to the frontal central P2 and P3 components.

**Figure 3.**
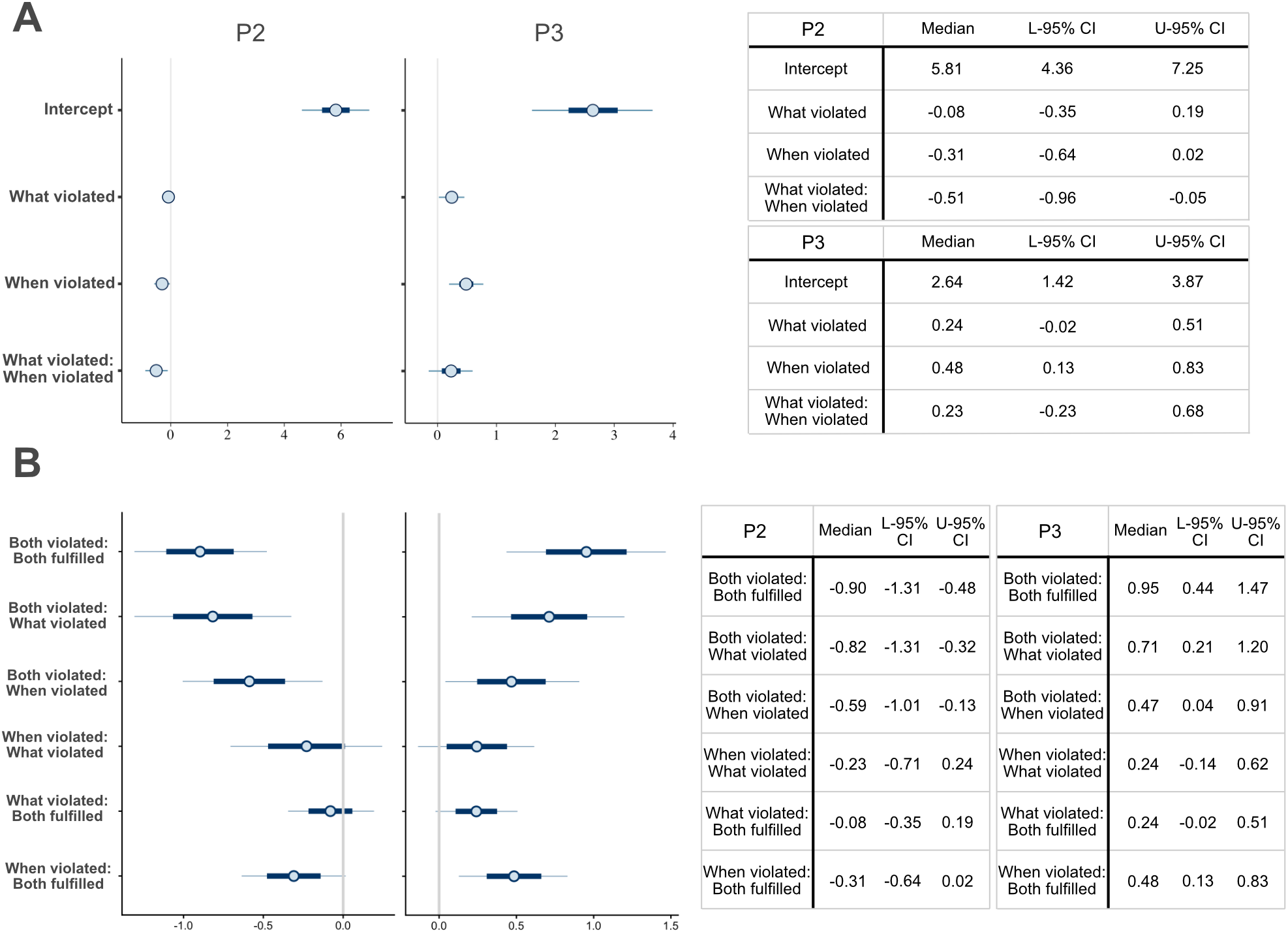
(A) Medians and credible intervals (On the plots: 50%, thick line; 90% thin line. On the tables: 95%) of parameter values in P2 and P3. Intervals that do not include zero have the denoted probability to be a true effect. (B). Medians and credible intervals (95%) of planned pairwise comparisons of condition estimates, calculated with the function ‘hypothesis ’in the R package ‘brms’. Intervals that do not include zero have the denoted probability to be a true effect, i.e., differences between the indicated factors to be significant.

### P2

The analysis of P2 amplitude revealed a significant interaction between “what” and “when” prediction violation (95% CI [–0.96, –0.05]), while no significant main effect was found for either the “what” or “when” violation alone (95% CI [–0.35, 0.19] and 95% CI [–0.64, 0.02] respectively). The planned comparisons showed that the what-violated and when-violated conditions were not significantly different from the both-fulfilled condition (95% CI [–0.35, 0.19] and 95% CI [–0.64, 0.02] respectively), and not significantly different from each other (95% CI [–0.71, 0.24]). Only the both-violated condition produced a significant effect, showing a reduction in P2 amplitude relative to the both-fulfilled condition (95% CI [–1.31, –0.48]). These results suggest that P2 amplitude reflects a conjunctive prediction, where a violation of either what or when prediction alone is insufficient to modulate the signal, and it is specifically the simultaneous violation of both that drives the P2 reduction.

### P3

The analysis of P3 amplitude showed a significant main effect for the when-violated condition (95% CI [0.13, 0.83]) but not for the what-violated condition (95% CI [–0.02, 0.51]), and no significant interaction between the two (95% CI [–0.23, 0.68]). The planned comparisons showed that the both-violated condition produced a significantly larger P3 amplitude compared to the both-fulfilled condition (+0.95 µV, 95% CI [0.44, 1.47]). The when-violated condition fell between these two, showing a P3 amplitude that was significantly larger than the both-fulfilled condition (+0.48 µV, 95% CI [0.13, 0.83]), yet significantly smaller than the both-violated condition (–0.47 µV, 95% CI [–0.91, –0.04]). Since the when-violated and both-violated conditions differ only in whether the what-prediction is also violated, the significant difference between them indicates that the violation of the what-prediction contributes an additional increment to the P3 response, despite showing no significant independent effect

Altogether, the results suggest that the PE response toward the mispredicted tone is driven by the co-occurrence of prediction violations associated with both action choice (what-prediction) and action timing (when-prediction).

## Discussion

The present study investigated whether predictions arising from two independent dimensions of voluntary action, namely what action to perform and when to perform it, are processed independently or integrated into a unified sensory expectation. We designed an active oddball paradigm in which participants learned independent associations between action choice and action timing and distinct auditory features, allowing violations of ‘what’ and ‘when’ predictions to occur independently or simultaneously within a fully crossed 2 x 2 design. Our ERP results revealed a dissociation between early and later stages of auditory processing. At the P2 component (∼150–180 ms), neither action-choice nor action-timing violations alone reliably modulated neural responses; instead, a significant reduction in amplitude emerged only when both ‘what’ and ‘when’ predictions were violated simultaneously, consistent with an early conjunctive representation of the expected action outcome. In contrast, the later P3 component (∼300–350 ms) exhibited a graded response, with action-timing violations producing a reliable increase in amplitude and additional modulation when both action dimensions were violated. Together, these findings suggest that the predictions derived from different dimensions of voluntary action were integrated during action-effect processing. They also indicate that the neural representation of this integration evolved over time. Specifically, the results are consistent with the idea that early sensory processing reflected a conjunctive, integrated representation of predicted action effects, whereas later stages involved a more graded evaluation of PE arising from both action dimensions. These results extend previous findings of action-effect prediction across isolated decision types (Chung et al., 2022) and complement recent evidence for the integration of perceptual ‘what’ and ‘when’ predictions (Cappotto et al., 2023), suggesting that the integration of multiple predictive signals may be a common computational principle across perception and action.

A key finding of our study concerns the pattern observed in the early P2 component: a significant reduction in amplitude was observed only when both the ‘what’ and ‘when’ predictions were violated simultaneously, whereas violations of either dimension alone did not produce a reliable modulation. This finding suggests that, at early stages of auditory processing (∼150–180 ms), the brain does not evaluate predictions from different action dimensions independently. Instead, it appears to treat the predicted sensory outcome as a unified, holistic template against which the actual sensory input is compared. Only when the incoming stimulus failed to match this integrated expectation on multiple dimensions did the system register a sufficiently large PE to modulate the P2 response.

Previous research has consistently linked the auditory P2 to the processing of action-effect PE, particularly in active oddball paradigms in which expected and unexpected consequences of voluntary actions are contrasted (Hsu, Hämäläinen, et al., 2015; Korka et al., 2019). Within predictive coding accounts, modulation of the P2 has been interpreted as reflecting the early comparison between predicted and incoming sensory information, with smaller responses to unexpected outcomes reflecting a mismatch between internal expectations and sensory input (Chung et al., 2022; Costa-Faidella et al., 2011; Grimm et al., 2011; Hsu, Hämäläinen, et al., 2015). Importantly, previous studies manipulated only a single source of prediction, such that the sensory outcome was predicted either by one action dimension or by a single learned action-effect association. Consequently, they could establish that PE arises when expected action consequences are violated, but not whether multiple action-derived predictions are represented independently or jointly.

Our findings suggest that, at least during this early stage of processing, predictions derived from action choice and action timing are integrated into a conjunctive representation of the expected sensory event. If the two prediction sources contributed independently to early sensory processing, isolated violations of either action dimension would have been expected to modulate the P2, with simultaneous violations producing approximately additive effects. Instead, neither isolated violation significantly altered P2 amplitude relative to the fully predicted condition, whereas simultaneous violations produced a robust modulation. This pattern is more consistent with the idea that the brain compares incoming sensory information against a unified representation of the expected action consequence than with independent comparisons of individual predicted stimulus features. In this respect, the present findings extend those of Chung et al. (2022). Whereas Chung and colleagues demonstrated that predictions can be generated independently from the ‘what’, ‘when’, and ‘whether’ components of intentional action, our results indicate that predictions arising from different intentional decisions do not simply coexist but are combined into a single expectation against which sensory consequences are evaluated.

Importantly, the absence of a significant P2 modulation following violations of a single action dimension does not imply that these violations go undetected. Rather, the present findings indicate that the early PE response indexed by the P2 does not scale independently with violations of action choice and action timing. Instead, P2 amplitude was modulated only when both predictions were violated simultaneously, suggesting that early PE processing is sensitive to the conjunction of multiple action-derived expectations rather than to violations of either prediction in isolation.

One possible interpretation is that the predictive system compares incoming sensory information against an integrated expectation of the anticipated action outcome. In the present task, action choice and action timing predicted different physical properties of the same auditory event. Rather than maintaining separate expectations for each predicted feature, the predictive system may combine these expectations into a unified prediction against which sensory input is evaluated. Under this account, violating a single predicted feature would preserve much of the expected sensory event, whereas violating both simultaneously would produce a substantially larger mismatch, resulting in the conjunctive P2 response observed here. Although the present data cannot determine the representational format underlying action-effect prediction, this interpretation is consistent with predictive coding theories proposing that hierarchical generative models predict coherent causes of sensory input rather than isolated stimulus attributes (Friston, 2005). It is also compatible with the ideomotor and event coding theories, which propose that voluntary actions are selected and represented in terms of their anticipated sensory consequences rather than isolated motor commands (Hommel, 2019; Prinz, 1997). Although these accounts do not explicitly address how multiple anticipated consequences are combined, the present findings suggest that this question merits further investigation.

The later P3 component revealed a markedly different pattern. Unlike the P2, P3 amplitude increased reliably following violations of action timing and showed an additional increment when action choice was also violated. Although action-choice violations alone did not significantly modulate the P3, the difference between the ‘when violated’ and ‘both violated’ conditions indicates that violations associated with action choice contributed to the later response once another prediction had already been violated. This graded pattern suggests that the processes indexed by the P3 differ qualitatively from those reflected in the earlier P2. Whereas the P2 appears to signal whether the predicted sensory event matches the integrated expectation, the P3 may reflect a subsequent stage in which the significance of the mismatch is evaluated, and the internal predictive model is updated. This interpretation accords with previous proposals linking the P3 to higher-order cognitive processes, including context updating, attentional allocation, and the behavioural evaluation of PE (Donchin & Coles, 1988; Polich, 2007). Unlike the earlier N1 and P2 ERP components, which are thought to reflect more automatic and feature-specific aspects of PE, the P3 is sensitive to the salience and task relevance of unexpected events (Friedman et al., 2001; Mars et al., 2008). From this perspective, the dissociation between P2 and P3 suggests that the integration of multiple action-derived predictions is not static but unfolds over time, with early sensory processing operating on a conjunctive representation and later processing reflecting a more graded evaluation in which information from different violated action dimensions contributes to the overall PE response and, potentially, to the updating of the internal predictive model (Friston, 2005; Garrido et al., 2009).

Building on this interpretation, the dissociation between the conjunctive P2 and the graded P3 suggests that the integration of action-derived predictions is dynamic rather than static. Whereas early sensory processing appears to operate on an integrated representation of multiple action-derived expectations, later processing reflects a more differentiated evaluation in which information form individual violated action dimensions contributes to the overall PE response. Thus, the present findings extend ideomotor and event coding accounts by suggesting not only that voluntary actions are represented in terms of their anticipated sensory consequences, but also that the neural evaluation of those anticipated consequences unfolds across successive stages of processing. Although the computational mechanisms underlying these stages remain to be established, this temporal progression is consistent with predictive coding accounts in which PEs evolve over successive stages of sensory and cognitive processing (Friston, 2005; Garrido et al., 2009).

The present findings also contribute to the broader literature on predictive processing by bridging action-based and perceptual prediction. A recent study (Cappotto et al., 2023) demonstrated that predictions about stimulus identity (“what”) and temporal structure (“when”) interact during passive auditory perception, such that PEs for sound identity depended on temporal expectations. Notably, this interaction emerged at an early latency (approximately 130–180 ms), comparable to the time window in which the conjunctive P2 effect was observed in the present study. Although perceptual predictions arise from statistical regularities in the external environment, whereas the predictions examined here were generated internally through voluntary actions and learned action-effect associations, both studies converge on the conclusion that content and temporal predictions are not evaluated independently during early sensory processing. While the present study was not designed to address the underlying neural circuitry, Cappotto and colleagues’ demonstration that these interactions involve distributed fronto-parietal networks raises the possibility that comparable integrative mechanisms may also contribute to action-effect prediction. Together, these findings suggest that the integration of content (“what”) and temporal (“when”) predictions may represent a general computational principle operating across both perceptual and motor-sensory domains.

Several limitations should be acknowledged. First, the present study examined only two dimensions of intentional action. Although Chung et al. (2022) demonstrated that ‘whether’ decisions can also generate action-effect predictions, it remains unknown how predictions arising from action execution interact with those derived from action choice and timing. Extending the present factorial approach to include the ‘whether’ dimension would provide a more complete account of predictive processing during intentional action. Second, the present paradigm employed auditory action effects exclusively. Whether similar integration mechanisms operate across other sensory modalities, or for multisensory action consequences, remains an important question for future research. Finally, although ERPs provide excellent temporal resolution for tracking PE processing, they offer only limited insight into the underlying neural circuitry. Combining similar paradigms with source localisation, magnetoencephalography, or functional neuroimaging may help determine whether action-based prediction integration recruits neural networks similar to those identified for perceptual prediction by Cappotto et al. (2023).

In conclusion, the present findings demonstrate that predictions derived from different dimensions of voluntary action are integrated during action-effect processing rather than being evaluated entirely independently. Early sensory processing was characterised by a conjunctive PE response that emerged only when multiple action-based expectations were violated simultaneously, whereas later processing reflected a more graded evaluation of individual prediction violations. Together with previous work on intentional action (Chung et al., 2022) and recent evidence from perceptual prediction (Cappotto et al., 2023), these findings suggest that the integration of “what” and “when” predictions is a general property of predictive processing that extends across both action and perception. More broadly, they indicate that predictive processing during voluntary action is dynamic, with the relationship between multiple action-derived predictions evolving over the course of sensory and cognitive evaluation.

## CRediT authorship contribution statement

**Wai Ying Chung:** Conceptualization, Formal analysis, Writing – original draft, Visualization, Methodology, Software, Visualization, Investigation. **Álvaro Darriba:** Conceptualization, Formal analysis, Writing – original draft. **Florian Waszak:** Conceptualization, Methodology, Supervision, Project administration, Funding acquisition, Writing – review & editing.

## Acknowledgements

This work was supported by the French Agence Nationale de la Recherche (ANR) within the *Programme franco-allemand en Sciences humains et sociales (FRAL*) 2016 (PROJECT ID: ANR-16-FRAL-0008 to F.W.) This research was partly supported by IdEx Université Paris Cité ANR-18-IDEX-0001.

